# IGD: A simple, efficient genotype data format

**DOI:** 10.1101/2025.02.05.636549

**Authors:** Drew DeHaas, Xinzhu Wei

## Abstract

**Motivation:** While there are a variety of file formats for storing reference-sequence-aligned genotype data, many are complex or inefficient. Programming language support for such formats is often limited. A file format that is simple to understand and implement – yet fast and small – is helpful for research on highly scalable bioinformatics.

**Results:** We present the Indexable Genotype Data (IGD) file format, a simple uncompressed binary format that can be more than 100 times faster and 3.5 times smaller than *vcf.gz* on Biobank-scale whole-genome sequence data. The implementation for reading and writing IGD in Python is under 350 lines of code, which reflects the simplicity of the format.

**Availability:** A C++ library reading and writing IGD, and tooling to convert .*vcf.gz* files, can be found at https://github.com/aprilweilab/picovcf. A Python library is at https://github.com/aprilweilab/pyigd

## Introduction

Genetic polymorphism data is typically stored in a tabular format that can be thought of as an *S* × *N* matrix. The rows represent *S* sites and the columns represent *N* individuals, where each site has at least one alternate allele that differs from the reference sequence, but may have many (a multi-allelic site). Variant Call Format (VCF) (Danecek *et al*., 2011) and its compressed form (.*vcf.gz*) are mainstays of tooling that process such tabular genotype data. The VCF format is very flexible, and its plaintext nature makes it easy to understand, construct, and parse. However, it is inefficient to store and process for large-scale datasets, as evidenced by the proliferation of faster and more compact formats. Among alternate formats, BCF (Li, 2011), BGEN (Band and Marchini, 2018), and BED (Purcell *et al*., 2007) have popular tooling support. Newer, more efficient formats such as PGEN (Rivas and Chang, 2024), XSI (Wertenbroek *et al*., 2022), Savvy (LeFaive *et al*., 2021), GTshark (Deorowicz and Danek, 2019), GRG (DeHaas *et al*., 2024), and others (Lan *et al*., 2021; Browning *et al*., 2018) leverage the similarity between samples at nearby genetic positions (due to linkage-disequilibrium (LD)) to compress genotype data to an impressive extent.

Here we present the Indexable Genotype Data (IGD) format, which is designed by the (sometimes at odds) principles of simplicity and efficiency. IGD encodes tabular genotype data as hard calls, similar to pVCF and BED. The only meta-data it stores (optionally) are identifiers for variants and individuals; the expectation is that most meta-data can be stored separately in general purpose file formats like CSV or JSON (Pezoa *et al*., 2016). IGD is uncompressed, which makes reading and writing the format easy to implement and avoids the need for external compression libraries which may not be easily usable across platforms or programming languages. IGD is a binary format, and supports multi-allelic variants, any ploidy up to 255, is contained in a single file, and can be constructed in one pass over the input data. IGD can represent both phased and unphased data, but all data in the file must have the same phasedness.

## Methods

Given that we have *N* individuals in a dataset, we number them *0…*(*N-1*). There are *N*_*H*_ = *N* × *ploidy* haploid samples of these individuals, which are similarly numbered *0…*(*N*_*H*_*-1*). Throughout when we refer to a “sample” we mean a *haploid* sample. There are *M* variants in a dataset, each of which can be uniquely identified by the pair (*base-pair position, alternate allele*), and *Q* variants that contain at least one sample with missing data.

There are a few significant aspects of the IGD format worth highlighting.

### All polymorphic sites are stored using bi-allelic format

Instead of storing a row per site (*S* × *N*_*H*_ matrix), IGD stores a row per variant (*M* × *N*_*H*_ matrix). Multi-allelic sites are supported by *expanding them* into a row per variant. For example, a (*k*+1)-allelic site with *k* alternate alleles (without missing data) is expanded into *k* variants that all have the same position and reference allele, but different alternate alleles. The original multi-allelic sites can be recovered by aggregating the IGD variants by position, however, keeping the data as an *M* × *N*_*H*_ matrix is often convenient for statistical genetics or population genetics applications.

In practice, IGD is an (*M* + *Q*) × *N*_*H*_ matrix, since missing data is encoded as a row of samples representing the ones with missing data, instead of representing those with the alternate allele.

### IGD contains an internal index

The index contains the genomic position (in base-pairs) of each variant, and can be cross-referenced to the genotype data, the allele strings, or variant IDs. By keeping the contents of the index small (16 bytes per variant) we keep the cost of reading it from disk very small. All variant-related data in IGD can be randomly accessed by *i*, the row number of that variant in the IGD index.

### IGD uses one of two compact genotype formats per variant

Each row of genotype data is represented as the set of samples that have the variant/alternate allele. The two simple, compact ways to store this data are either (a) sparsely as a list of sample numbers or (b) densely as a bit-vector where each bit at position *i* reflects whether the *i*-th sample has the alternate allele (1) or not (0). We can choose between representation (a) and (b) by examining the allele frequency *p*_*i*_ of the variant in question. Sample numbers are represented by 32-bit unsigned integers, which means that if 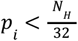 the sparse representation is more compact (a), otherwise the bit-vector representation (b) is smaller. It is important to note that a valid IGD file can be constructed using *only* row representation (a) or *only* bit vector representation (b), or any mixture of the two, but the optimally sized IGD will determine which to use on a per-variant basis.

Converting between these two representations is trivial, and thus the representation on disk does not have to be the representation used for computation.

### 2.1 File Format Details

The layout of an IGD file is shown in **Fig. 1**. The header contains file offsets for each of the sections after the genotype data, as their positions are unpredictable otherwise, and random access to them is useful. We use the following storage type definitions.

**Fig. 1.**
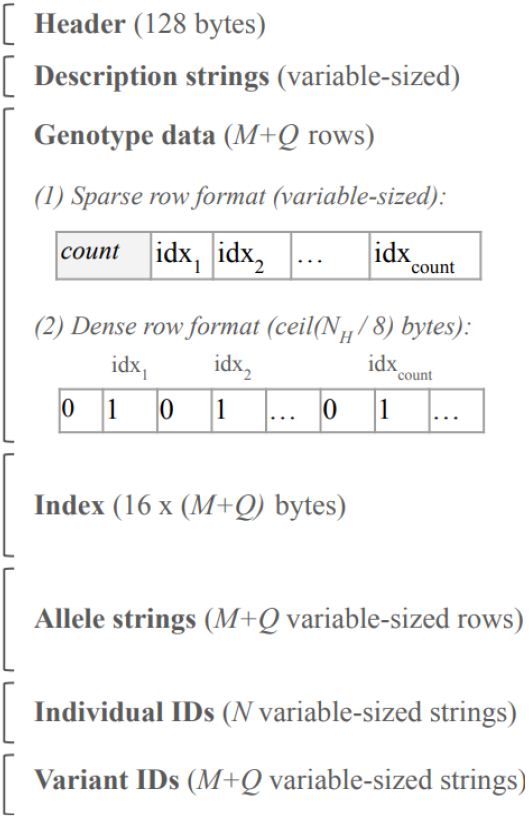
IGD file layout. The layout of an IGD file on disk. *M* is the number of variants, *N* is the number of individuals, and *N*_*H*_ is the number of haploid samples. Genotype data rows may be either sparse (a list of haploid sample indexes containing the alternate allele) or dense (a bit-vector with a 1 at an index *i* iff the *i*^th^ haploid sample contains the alternate allele).

***uint32*:** A 32-bit unsigned integer.

***uint64*:** A 64-bit unsigned integer.

***string*:** A uint32 for the length *k*, followed by *k* bytes for the contents.

***list32*:** A uint32 for the length *k*, followed by *k* uint32 values for the contents.

***bv(w)*:** A bit-vector of *w* bits stored at the byte granularity. The number of bytes used is *ceil*(*w*/8). Given a sample index *b* that we want to store as a 1, the byte offset is determined by *floor*(*b*/8). Within that byte we set the (*7 -* (*b mod 8*))^th^ least significant bit; that is, if (*b mod 8*) *= 0* we will set the most significant bit.

### 2.1.1 Header

The header is a fixed-size (128 byte) table as described in Table 1.

**Table 1.** IGD header details.

| Byte offset | Datatype | Description |
| --- | --- | --- |
| 0 | uint64 | Magic number 0x3a0c6fd7945a3481 |
| 8 | uint64 | File format version |
| 16 | uint32 | Ploidy |
| 20 | uint32 | Sparsity threshold |
| 24 | uint64 | Number of variants |
| 32 | uint32 | Number of individuals $N$ |
| 36 | uint32 | Reserved for future use |
| 40 | uint64 | 64 bits of flags. |
| 48 | uint64 | File offset where the **Index** is located |
| 56 | uint64 | File offset where the **Allele strings** are located |
| 64 | uint64 | File offset where the **Individual IDs** are located |
| 72 | uint64 | File offset where the **Variant IDs** are located |
| 80 | 48 bytes | Reserved for future use |

### 2.1.2 Flags

The least-significant bit of the flags (in the header) signifies phasedness, where a value of 1 indicates phased data.

#### 2.1.2 Description strings

Immediately following the header are two *string* values. The first is a *string* describing how the file was created (e.g., “converted from foo.vcf.gz”) and the second is a generic description field.

#### 2.1.3 Genotype data

Immediately following the description strings is *M+Q* rows of genotype data, where each row is either a ***list32*** or a ***bv*(*N***_***H***_). The Index (described next) has a flag that indicates the type of each row.

#### 2.1.4 Index

The Index is *M+Q* rows of 16 bytes each, and can be viewed as two *uint64* values. The first ***uint64*** value contains the base-pair position associated with the variant in the least-significant 48 bits, followed by an 8-bit unsigned integer *numCopies*, and finally bitwise flags in the most-significant 8 bits. The currently defined flags are:

- **SPARSE=0×01**: If this flag is set the corresponding genotype row is a *list32*, otherwise it is a ***bv*(*N***_***H***_).
- **IS_MISSING=0×02**: If this flag is set the corresponding genotype row’s sample list represents missing data. The list of samples in the row do not have a variant call for that polymorphic site.

The *numCopies* value is 0 for phased data, but is 1≤ *numCopies* ≤ *ploidy* for unphased data.

The second ***uint64*** value in the Index row contains the file offset of the genotype row for the current variant. The *i*^th^ variant can be randomly accessed directly at *IndexStart* + (16 × *i*).

#### 2.1.5 Allele strings

This section is *M+Q* rows, each row contains first the ***string*** for the reference allele and then the ***string*** for the alternate allele.

#### 2.1.6 Individual IDs

This section is a ***uint64*** for the number of strings, followed by that many ***strings***, where the *k*^th^ is the identifier for individual *k*. This section is only present in the file if the corresponding header file offset entry is non-zero.

#### 2.1.7 Variant IDs

This section is a ***uint64*** for the number of strings, followed by that many ***strings***, where the *i*^th^ one is the variant identifier for the *i*^th^ variant. This section is only present in the file if the corresponding header file offset entry is non-zero.

### 2.2 Phasedness

Unphased data is stored by clearing the phased flag in the header, and storing separate variants for each number of copies of each alternate allele. The *numCopies* value in the index (see above) indicates the zygosity of the currently stored sample list. Additionally, instead of storing haploid sample lists, IGD stores individual-based sample lists for unphased data. That is, it represents an (*M* + *Q*) × *N* (instead of *N*_*H*_). matrix

### 2.3 Access Patterns

There are two typical access patterns for an IGD file. If the variant index *i* is known, we can seek directly to it in the Index and then seek directly to the genotype data for that variant. If any of the string data is needed (alleles, individual IDs, variant IDs) those tables will need to be loaded into memory so they are indexable by *i*, or just scanned on disk to find the *i*^th^ entry.

Alternatively, traversing an IGD file is done by seeking to the start of the Index. Starting at variant *i* = 0, each row of the Index is read and if the base-pair position is of interest then the genotype data is accessed using the file offset found in the current (*i*^th^) row of the Index. String data can be read into RAM one time, or a file pointer can be maintained to the current entry for each string table and incremented whenever *i* is incremented.

## Results

We compared file size and data traversal time between IGD, .*vcf.gz*, BCF, and PGEN (**Fig. 2**). These formats were chosen for their apparent popularity as well as for capturing a spectrum from simple and inefficient (.*vcf.gz*) to more complex, yet very efficient (PGEN). PGEN is on the more complex side because it uses LD-based compression, and also supports 8 different storage modes, some of which are for backwards compatibility (Chang, 2024).

**Fig. 2.**
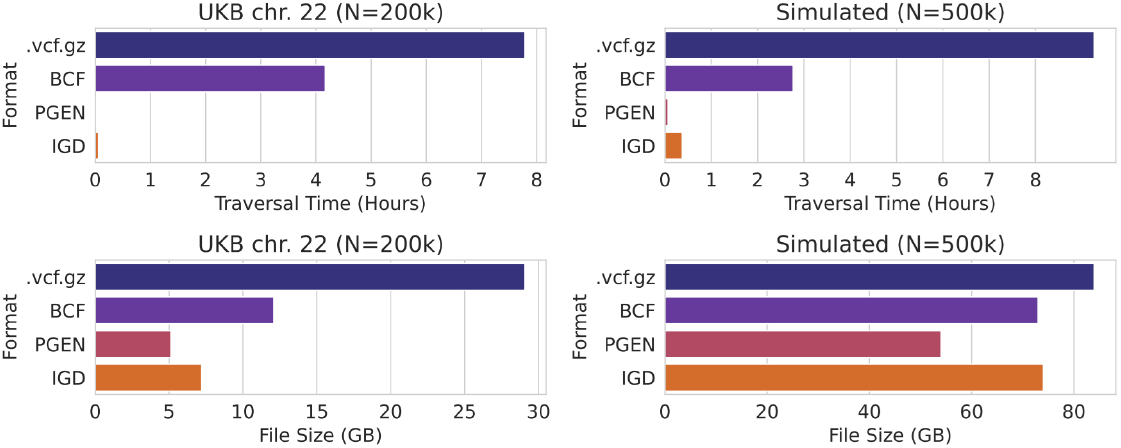
Comparison of file formats. The upper panels show time to traverse the file for UK Biobank WGS data (left) and simulated data (right). The lower panels show the file sizes for UK Biobank WGS data (left) and simulated data (right).

Allele frequency calculation was used for traversal time, as it is a trivial calculation over all the data in the file. For formats that encode an allele count, such as BCF, PGEN, and IGD (for sparse variants), we do not use that count but read the full sample data for each variant in order to measure the data traversal overhead. plink2 (Purcell and Chang, 2024)) was used for conversion to BCF and PGEN formats, as well as for allele frequency calculation for .*vcf.gz* and BCF. The API *PgrGetDifflistOrGenovec()* was used for calculating PGEN allele frequency. The BCF files were stripped of additional meta-data, and only contained variants and genotypes.

The simulated data was generated via stdpopsim (Adrion *et al*., 2020) and msprime (Kelleher *et al*., 2016), using an out-of-Africa demographic model (Jouganous *et al*., 2017) and sampling 500,000 European individuals, in an attempt to generate data similar to the UK Biobank whole-genome sequence (WGS) data (Hofmeister *et al*., 2023).

File sizes are all fairly similar on the simulated dataset, ranging from 54GB (PGEN) to 84GB (.vcf.gz), with IGD and BCF being similarly sized at 73-74GB. PGEN and IGD are the two smallest formats on the UKB dataset, which is much richer in low-frequency variants (∼96% variants are MAF<0.1% (Hofmeister *et al*., 2023)) than our simulated dataset (∼73% of variants are MAF<0.1%), and thus more compactly stored with a sparse representation.

BCF and .*vcf.gz* rely heavily on standard compression algorithms, which is illustrated by the traversal times for IGD and PGEN being many times faster, especially on the UKB data. PGEN is the smallest and fastest file format, about 30% smaller and 5x faster than IGD.

File format conversion times are summarized in **Table 2**. PGEN is again fastest, likely partially due to plink’s highly optimized .*vcf.gz* read functionality.

**Table 2.** Conversion times from .*vcf.gz*.

| Dataset | File Format | Conversion time (hours) |
| --- | --- | --- |
| UKB chr. 22 (N=200k) | BCF | 32.5 |
| UKB chr. 22 (N=200k) | PGEN | 13.6 |
| UKB chr. 22 (N=200k) | IGD | 21.3 |
| Simulated (N=500k) | BCF | 29.9 |
| Simulated (N=500k) | PGEN | 9.3 |
| Simulated (N=500k) | IGD | 23.6 |

The compactness and simplicity of IGD format make it easily usable in bioinformatic tool development. IGD has been successfully used as the input to Genotype Representation Graph (GRG) construction (DeHaas *et al*., 2024) construction, and is essential for the efficiency of that process on biobank-scale data. GRG construction requires fast indexing of genomic regions as well as fast genotype data access. Indexing compressed formats (such as .vcf.gz and BCF) can be complex, and creates a separate file for the index. For IGD the index is a fundamental part of the file format. **Fig. 3** shows the time to construct a GRG tree, the first part of GRG construction, is 13-15x faster for IGD than for .*vcf.gz*. We extracted the same region (for varying lengths, on the x-axis) from a simulated dataset with 1 million haploid samples, and then timed the GRG tree construction for that region.

**Fig. 3.**
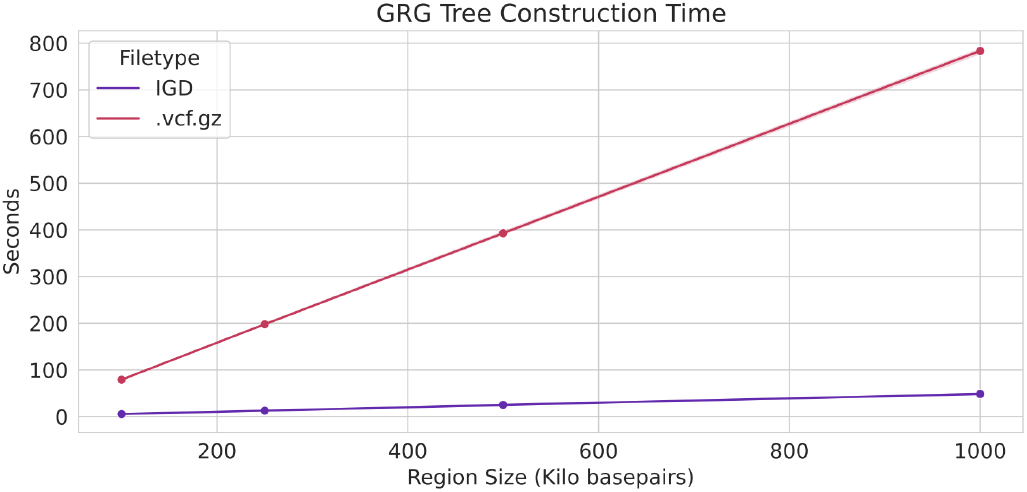
File format impact on GRG tree construction. Genotype Representation Graph (GRG) tree construction time for .vcf.gz vs. IGD file formats, for small regions of the genome (x-axis) from a simulated dataset with 1 million haploid samples.

IGD provides an extremely simple, yet efficient, alternative to existing file formats. It focuses on genotype data storage, and ease-of-use for developers of scalable prototypes and tools.

## Acknowledgements

This research has been conducted using the UK Biobank Resource under Application Number 97908.

